# Immunopeptidome profiling of human coronavirus OC43-infected cells identifies CD4 T cell epitopes specific to seasonal coronaviruses or cross-reactive with SARS-CoV-2

**DOI:** 10.1101/2022.12.01.518643

**Authors:** Aniuska Becerra-Artiles, Padma P. Nanaware, Khaja Muneeruddin, Grant C. Weaver, Scott A. Shaffer, J. Mauricio Calvo-Calle, Lawrence J. Stern

**Affiliations:** Department of Pathology, Department of Biochemistry and Molecular Biotechnology, UMass Chan Medical School, Worcester MA; Mass Spectrometry Facility, UMass Chan Medical School, Shrewsbury MA; Department of Biochemistry and Molecular Biotechnology, UMass Chan Medical School, Worcester, MA 01655, USA

## Abstract

Seasonal “common-cold” human coronaviruses are widely spread throughout the world and are mainly associated with mild upper respiratory tract infections. The emergence of highly pathogenic coronaviruses MERS-CoV, SARS-CoV, and most recently SARS-CoV-2 has prompted increased attention to coronavirus biology and immunopathology, but identification and characterization of the T cell response to seasonal human coronaviruses remain largely uncharacterized. Here we report the repertoire of viral peptides that are naturally processed and presented upon infection of a model cell line with seasonal human coronavirus OC43. We identified MHC-I and MHC-II bound peptides derived from the viral spike, nucleocapsid, hemagglutinin-esterase, 3C-like proteinase, and envelope proteins. Only three MHC-I bound OC43-derived peptides were observed, possibly due to the potent MHC-I downregulation induced by OC43 infection. By contrast, 80 MHC-II bound peptides corresponding to 14 distinct OC43-derived epitopes were identified, including many at very high abundance within the overall MHC-II peptidome. These peptides elicited low-abundance recall T cell responses in most donors tested. In vitro assays confirmed that the peptides were recognized by CD4+ T cells and identified the presenting HLA alleles. T cell responses cross-reactive between OC43, SARS-CoV-2, and the other seasonal coronaviruses were confirmed in samples of peripheral blood and peptide-expanded T cell lines. Among the validated epitopes, S_903-917_ presented by DPA1*01:03/DPB1*04:01 and S_1085-1099_ presented by DRB1*15:01 shared substantial homology to other human coronaviruses, including SARS-CoV-2, and were targeted by cross-reactive CD4 T cells. N_54-68_ and HE_128-142_ presented by DRB1*15:01 and HE_259-273_ presented by DPA1*01:03/DPB1*04:01 are immunodominant epitopes with low coronavirus homology that are not cross-reactive with SARS-CoV-2. Overall, the set of naturally processed and presented OC43 epitopes comprise both OC43-specific and human coronavirus cross-reactive epitopes, which can be used to follow T cell cross-reactivity after infection or vaccination and could aid in the selection of epitopes for inclusion in pan-coronavirus vaccines.

**Author Summary:** There is much current interest in cellular immune responses to seasonal common-cold coronaviruses because of their possible role in mediating protection against SARS-CoV-2 infection or pathology. However, identification of relevant T cell epitopes and systematic studies of the T cell responses responding to these viruses are scarce. We conducted a study to identify naturally processed and presented MHC-I and MHC-II epitopes from human cells infected with the seasonal coronavirus HCoV-OC43, and to characterize the T cell responses associated with these epitopes. We found epitopes specific to the seasonal coronaviruses, as well as epitopes cross-reactive between HCoV-OC43 and SARS-CoV-2. These epitopes should be useful in following immune responses to seasonal coronaviruses and identifying their roles in COVID-19 vaccination, infection, and pathogenesis.

## Introduction

Coronaviruses are single-stranded RNA viruses of the genus *Nidovirales*, family *Coronaviridae* that infect vertebrates. Seven species in the *Orthocoronavirinae* sub-family are known to infect humans, with a wide range of pathogenicity [1]. Human coronavirus (HCoV) 229E and NL63 in the *alpha-coronavirus* genus, and OC43 and HKU1 in the *beta-coronavirus* genus, are associated with mild upper-respiratory-tract infections and common colds. In contrast, SARS-CoV, MERS-CoV, and SARS-CoV-2, all in the *beta-coronavirus* genus, are associated with a severe respiratory syndrome [2]. Common-cold-associated seasonal HCoVs are widespread and infect humans in seasonal waves [3–5]. OC43 is closely related to bovine coronavirus (BCoV) and was initially isolated in 1967 from individuals with upper respiratory tract infections [6]. Among the seasonal human coronaviruses, OC43 is believed to have emerged most recently. Molecular clock analysis of the spike gene sequences suggests a relatively recent zoonotic transmission event and dates their most recent common ancestor between 1890 to 1923 [7–9]. This led to the proposal that the 1898 pandemic (“Russian Flu”), which caused a worldwide multi-wave outbreak killing preferentially older individuals similar to COVID-19, may have been the result of the emergence of OC43 [10]. The OC43 reference genome (ATCC-VR-759) spans 30,738 kbp, encoding 10 ORFs which are translated into 24 proteins [11].

Before the emergence of the pandemic coronavirus SARS-CoV-2, few studies characterized the immune response to the seasonal HCoVs, which account for ∼10-30% of common colds [12,13]. Studies of T cell responses to HCoVs and the identification of epitopes driving them are scarce. Before the SARS-CoV-2 pandemic, Nilges et al. identified a coronavirus MHC-I epitope derived from the OC43 NS2 protein using MHC-binding prediction algorithms and showed that T cell responses were cross-reactive with a human papillomavirus 16 epitope [14]. Later, Boucher et al. studied T cell responses to OC43 and 229E viral antigens and to multiple sclerosis (MS) autoantigens in MS patients. Virus-specific T cell clones were isolated, including 34 clones responding to OC43, as well as 10 T cell clones cross-reactive with HCoV and MS autoantigens, but the specific viral epitopes were not identified [15]. More recently, after the rise of SARS-CoV-2, Woldemeskel et al [16] reported T cell responses to pools of spike, nucleoprotein, and membrane proteins of the four seasonal coronaviruses. Peptide responses to the spike protein of NL63 were deconvoluted resulting in the identification of 22 target peptides, of which 3 are SARS-CoV-2 cross-reactive and the remaining 19 are HCoV-specific novel epitopes.

Studies of T-cell responses to SARS-CoV and SARS-CoV-2 have reported that responding T cell populations are present in blood samples collected before the emergence of these viruses [17,18]. This led to the suggestion that pre-existing immunity, potentially elicited by a previous infection(s) with seasonal HCoVs, could be responsible for these responses, and prompted a search for the cross-reactive epitopes responsible. In fact, most OC43 epitopes reported in the Immune Epitope Database [19] were identified in the context of HCoV/SARS-CoV-2 cross-reactivity studies. Schmidt et al. used a highly conserved peptide derived from the SARS-CoV-2 nucleoprotein to identify cross-reactive MHC-I responses and found that homologous HCoV peptides, including one from OC43, also were recognized [20]. Mateus et al. used overlapping SARS-CoV-2 peptides to screen for cross-reactive responses in unexposed donors and identified six MHC-II epitopes from five source proteins, for which responses to the OC43 homologs could also be observed [21]. Keller et al. identified a cross-reactive OC43 epitope derived from the nucleocapsid, which induced responses in SARS-CoV-2 specific T cells expanded from COVID-19 recovered donors using SARS-CoV-2 antigens [22]. Ferretti et al. reported a cross-reactive MHC-I epitope derived from the nucleocapsid, highly conserved among beta-coronaviruses [23], and Lineburg et al. found that the immunodominant response to this peptide is widespread in HLA-B7+ individuals, both recovered COVID-19 and unexposed [24]. Our previous work [25] and other studies [21,26–30] identified a highly conserved and cross-reactive MHC-II SARS-CoV-2 epitope (S_811-831_), derived from a conserved region in the spike protein and presented in the context of HLA-DP4 (DPA1*01:03/DPB1*04:01), HLA-DP2 (DPA1*01:03/DPB1*02:01), and HLA-DQ5 (DQA1*01:01/DQB1*05:01) [25]. Despite these advances, an unbiased approach to the identification of OC43 T cell epitopes independent of SARS-CoV-2 reactivity has not been reported.

T cell epitope identification can be approached in different ways, including screening of overlapping peptide libraries, predicting potential epitopes using MHC-binding prediction algorithms, or identifying naturally processed and presented peptides eluted from purified MHC molecules isolated from infected cells. In this work, we used the latter method, which has proven to be very efficient in identifying immunogenic peptides in human T cell responses to vaccinia virus [31–33], HHV-6B [34], influenza [35], measles [36], EBV [37], and SARS-CoV-2 [38] and in mouse responses to vaccinia virus where this was validated extensively [39]. Here, we identified and characterized naturally-processed viral epitopes presented by HEK293 cells transfected with master transcriptional regulator CIITA and infected with OC43. CIITA served to upregulate the expression of MHC-II molecules and associated antigen presentation machinery (reviewed in [40]), as in previous studies [34,41–43]. Overall, 83 naturally processed viral peptides were identified: 3 peptides were identified as associated with the MHC-I proteins HLA-A*02:01 or HLA-B*07:02, and 80 viral peptides representing length variants of 14 unique MHC-II epitopes were identified as associated with HLA-DRB1*15:01, HLA-DRB5*01:01, or HLA-DPA1*01:03/DPB1*04:02. T cell responses to 11 of the peptides were observed in partially HLA-matched donors, confirming the immunogenicity of these peptides. Among the naturally presented peptides identified was S_901-920_, orthologous to a highly conserved, frequently identified, cross-reactive SARS-CoV-2 epitope S_811-831_.

## Results

### Characterization of MHC-I and MHC-II immunopeptidomes presented in OC43-infected cells

Our experimental approach to the identification and characterization of naturally processed epitopes is diagrammed in Fig 1A. Peptide-MHC complexes carrying naturally processed and presented peptides were isolated by immunoaffinity from OC43-infected cells, and bound peptides were eluted and characterized by mass spectrometry. Next, peptides corresponding to the naturally processed epitopes were synthesized, tested for HLA binding, and used for evaluation of T cell responses in mononuclear cells from peripheral blood samples. We used the cell line HEK293, which is homozygous in all MHC-I and MHC-II loci and susceptible to being infected with the OC43 virus. The HLA alleles present in this cell line are: A*02:01 (A2), B*07:02 (B7), C*07:02 (C7), DRB1*15:01 (DR2b), DRB5*01:01 (DR2a), DPA1*01:03/DPB1*04:02 (DP4.2), and DQA1*01:02/DQB1*06:02 (DQ6.2).

**Figure 1.**
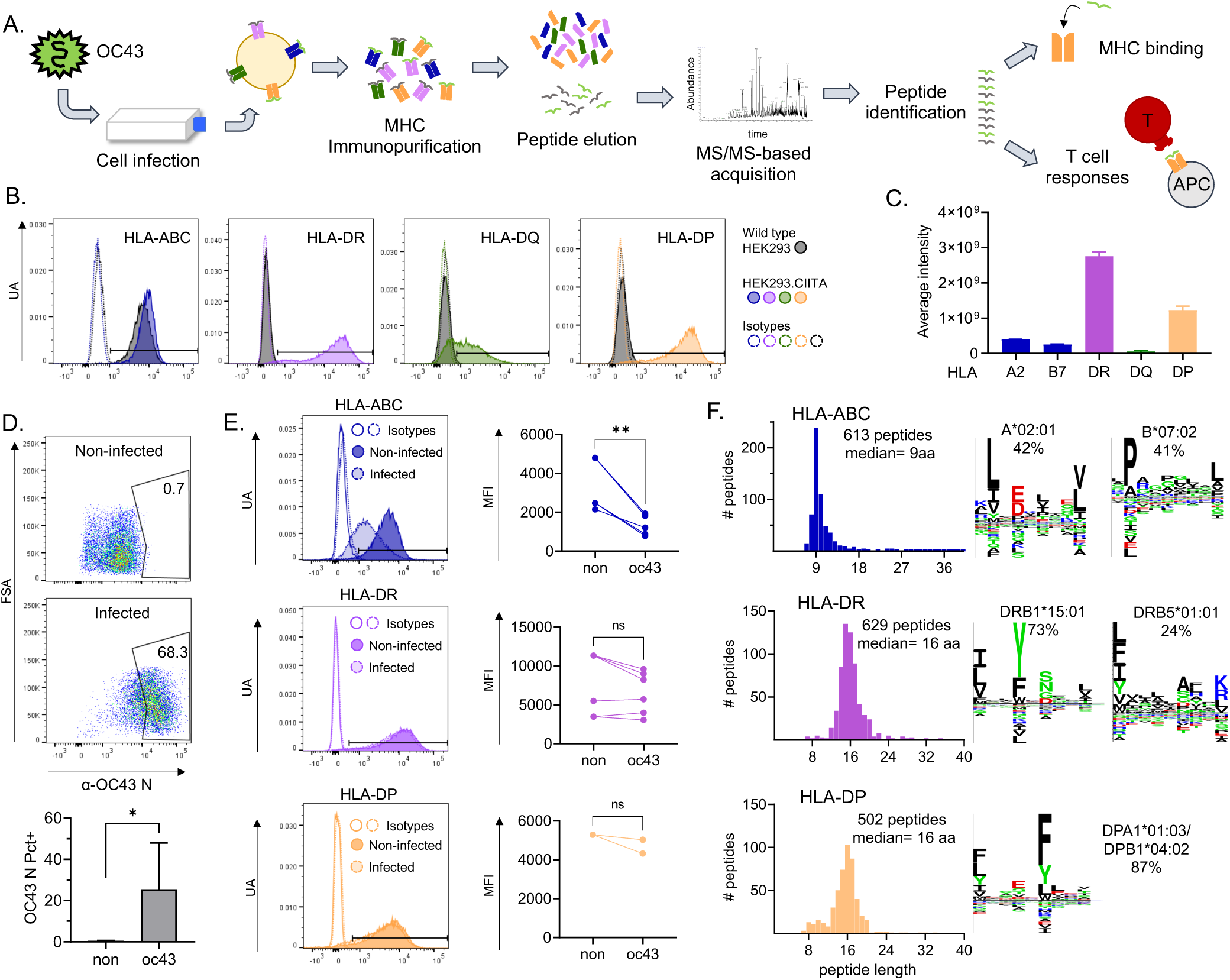
Immunopeptidome workflow and HLA-ABC, HLA-DR, and HLA-DP immunopeptidomes in OC43-infected HEK293 cells. **A.** Experimental approach: HEK293 cells transduced with CIITA were infected with OC43. After 3 days, cells were collected and pMHC complexes were purified by immunoaffinity. Peptides were eluted from pMHC and analyzed by LC-MS/MS for identification. Identified peptides were used in biochemical and immunological assays. **B.** MHC expression on the surface of HEK293 cells. Four panels corresponding to the surface expression of HLA-ABC, HLA-DR, HLA-DQ, and HLA-DP are shown. HLA levels on wild-type cells are shown by grey histograms. HLA levels after transduction with CIITA are shown by colored histograms: HLA-ABC (blue), HLA-DR (purple), HLA-DQ (green), and HLA-DP (yellow). Isotype control staining is shown as an open histogram with dotted lines, following the same color scheme. **C.** Levels of total HLA-DR, HLA-DP, and HLA-DQ proteins in CIITA-transfected HEK293 cells measured by label-free quantitative proteomics. **D.** Representative dot plots of intracellular staining for OC43 nucleoprotein in non-infected cells (top) and at 3 days after infection (bottom). **E.** Representative histograms showing the comparison of surface levels of HLA-ABC, HLA-DR, and HLA-DP on non-infected (dark histograms) and infected (light histograms) cells. Graphs show the MFI in non-infected (non) and infected (oc43) cells from 3-6 independent infections. Statistical analysis in D and E by paired t-test, * p<0.05, ** p<0.01, ns: not significant. **F.** Length distribution of HLA-ABC, HLA-DR, and HLA-DP eluted immunopeptidomes (histograms). **G.** Sequence logos of clusters obtained using the Gibbs clustering analysis of HLA-ABC, HLA-DR, and HLA-DP eluted immunopeptidomes; percentage of peptides in each cluster and probable allele are shown.

We measured the expression of MHC-I and MHC-II on the surface of HEK293 cells using antibodies recognizing the three MHC-I proteins HLA-ABC or the individual MHC-II proteins HLA-DR, HLA-DQ, and HLA-DP. Expression of HLA-ABC was detected, but levels of HLA-DR, HLA-DP, and HLA-DQ were very low or below detection limits (Fig 1B, wild type HEK293). To induce expression of MHC-II, HEK293 cells were transduced with CIITA, the MHC-II master transcriptional regulator that controls the expression of MHC-II genes along with MHC-II processing and editing factors such as HLA-DM and cathepsins (reviewed in[40]). Transduced cells successfully upregulated the expression of HLA-DR and DP (Fig 1B, HEK293.CIITA), although HLA-DQ levels remained low. To confirm the low HLA-DQ expression level, the relative amounts of total MHC proteins were measured using a quantitative proteomics analysis (Fig 1C, Table S1). The levels of HLA-DQ were ∼20-fold lower than HLA-DR and HLA-DP. Thus, we restricted immunopeptidome analysis to HLA-ABC, HLA-DR, and HLA-DP.

HEK293.CIITA cells were infected with OC43 strain VR-759 at a multiplicity of infection of 0.1 and harvested on day 3 post-infection. Intracellular staining for OC43 nucleoprotein (N) showed a clear positive population of virus-infected cells at harvest, as compared to non-infected cells (Fig 1D). In 6 biological replicates, we observed that 11-68% of the HEK293.CIITA cells were positive for OC43 nucleoprotein expression.

Viruses have evolved many mechanisms to evade the immune system, including the downregulation of MHC proteins [44–46]. To assess the effect of OC43 infection on the expression of MHC-I and II on HEK293.CIITA cells, we evaluated the surface expression of HLA-ABC, HLA-DR, and HLA-DP after infection. The levels of HLA-ABC were significantly reduced after infection (an average of 60% reduction in median fluorescence intensity (MFI)), while the expression of HLA-DR and HLA-DP were mostly not affected (less than 10% reduction in MFI) (Fig 1E). This suggests that OC43 has a specific effect on the expression of MHC-I. While no apparent effect was observed for MHC-II, it is possible that CIITA transfection counteracts any effect of virus infection in our system as reported for SARS-CoV-2 and Ebola viruses [47].

We used a conventional immunoaffinity peptidomics workflow to identify peptides presented by MHC molecules in the infected cells. We purified MHC-bound complexes of two independent infections (62 and 116 x10^6^ cells) using immunoprecipitation after detergent solubilization of the membrane fraction of OC43-infected HEK293.CIITA cells. We used sequential immunoaffinity purification with anti-HLA-DR (LB3.1), anti-HLA-DP (B7/21), and anti-HLA-ABC (W6/32) antibodies, collecting three immunoprecipitated samples, one from each antibody, per biological replicate infection. The MHC-bound peptides were released from the purified MHC complexes by acid treatment, separated from the MHC protein subunits, and the resulting peptide mix was analyzed by LC-MS/MS for sequence identification. A database containing human and OC43 protein sequences was used for peptide assignment, with false-discovery rate (FDR) of 4.2%. The total immunopeptidome of infected cells consisted of 1,744 unique peptides (613 HLA-ABC, 629 HLA-DR, and 502 HLA-DP, Table S2a-c). The eluted peptides showed the expected length distribution peaking at 9 aa for HLA-ABC and 15-16 aa for HLA-DR and -DP, although HLA-DP showed a small peak of 8-11 residue peptides that might include non-specifically bound species [48] (Fig 1F). The immunopeptidome comprises both viral and host protein-derived peptides, with ∼96% of host-derived peptides.

The eluted peptide pools contain contributions from multiple MHC proteins. HLA-ABC eluted peptides were a mix of peptides eluted from the three MHC-I proteins present in HEK293.CIITA. cells. Likewise, HLA-DR peptides were a mix of peptides eluted from the genetically-linked DRB1*15:01 and DRB5*01:01 proteins. To help deconvolute these mixtures of peptides, we used unsupervised Gibbs clustering [49] of the eluted sequences in each sample. This analysis showed the presence of 2 motifs for MHC-I, representing 42 and 41% of the sequences (Fig 1F, HLA-ABC). These motifs closely matched those previously characterized for A*02:01 and B*07:02 by NetMHCpan [50,51], as shown in Figure S1. The characteristic C*07:02 motif [50,52] was not observed in the clustering analysis. We observed 2 motifs for HLA-DR, representing 73 and 24% of the sequences (Fig 1F, HLA-DR). The more abundant motif closely matched that previously characterized for DRB1*15:01 (DR2b), and the less abundant motif matched that for DRB5*01:01 (DR2a) [50,53] (Figure S1). For HLA-DP, one motif representing 87% of the sequences was observed (Fig 1F, HLA-DP), closely matching the expected DPA1*01:03/DPB1*04:02 motif [50,54] (Figure S1). Sequences not present in these clusters could represent non-canonical binders, ambiguities in the clustering for motif analyses, or the presence of non-specific peptides. For each eluted peptide, binding predictions for the relevant MHC-I (NetMHCpan 4.1), or MHC-II (NetMHCIIpan 4.0) proteins are shown in Table S2a-c. For DP4 we include predictions for both DPA1*01:02/DPB1*04:01 (DP4.1) and DPA1*01:02/DPB1*04:02 (DP4.2); peptides were eluted from DP4.2 cells, but the closely related DP4.1 protein was used for MHC-peptide binding studies and both DP4.1 and/or DP4.2 expressing donors were used for T cell studies (see below).

### Identification of viral peptides presented by HLA-ABC, HLA-DR, and HLA-DP

Within the immunopeptidome eluted from OC43-infected HEK293.CIITA cells, a total of 83 peptides corresponded to sequences from the OC43 virus (Table S2d). Among the viral peptides, 3 were eluted from HLA-ABC, 35 from HLA-DR, and 45 from HLA-DP, representing 0.6, 5.6, and 9.4% of the peptides isolated from each type of MHC protein. The average length of the viral peptides was consistent with that observed for the total peptides, with a peak at 9 residues for the MHC-I and around 15-16 residues for the MHC-II (Fig 2A).

**Figure 2:**
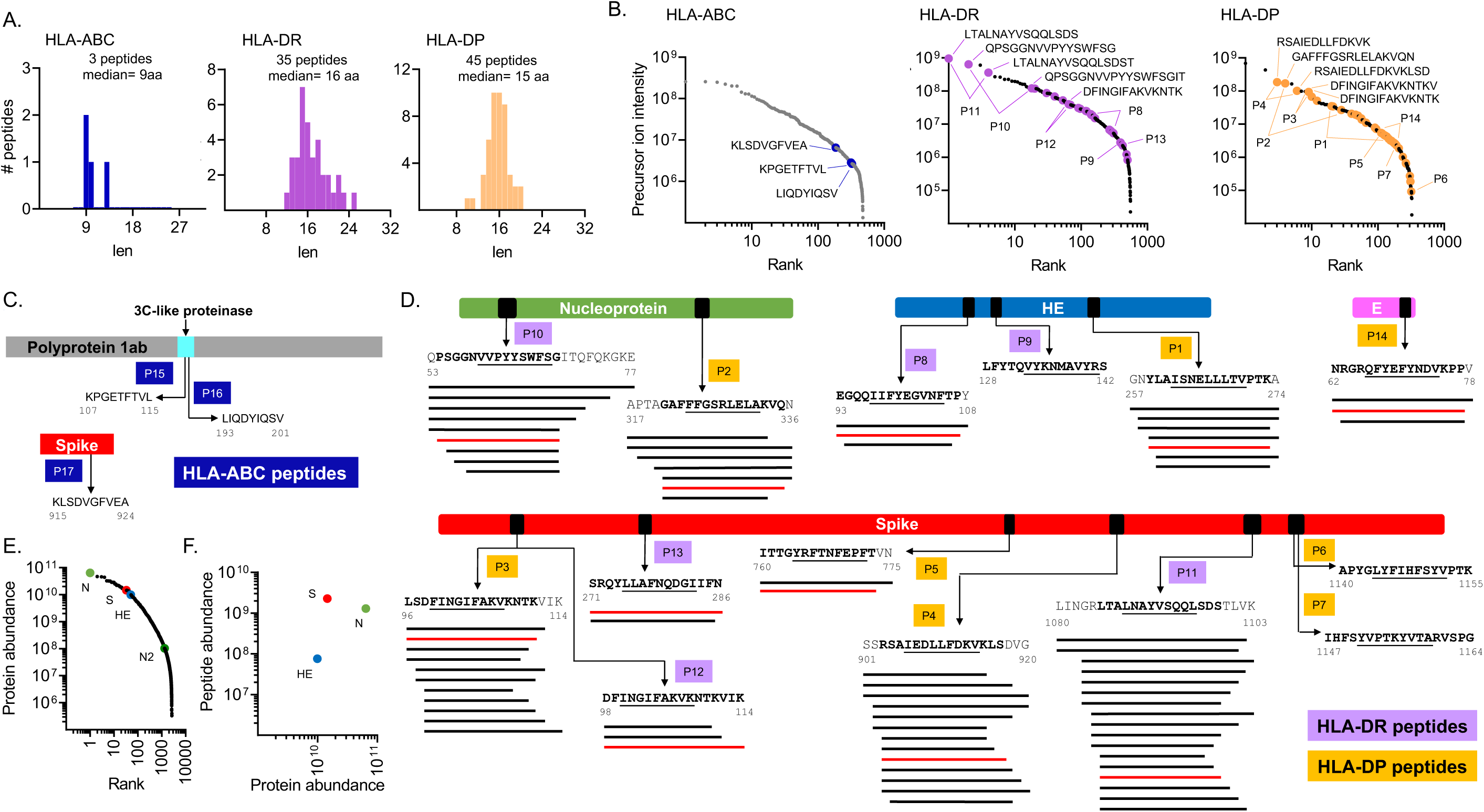
OC43 virus-derived peptides in the HLA-ABC, HLA-DR, and HLA-DP immunopeptidomes. **A.** Length distribution of virus-derived peptides within the HLA-ABC, HLA-DR, and HLA-DP immunopeptidomes of OC43-infected cells. **B.** Ranking of all HLA-ABC, HLA-DR, and HLA-DP eluted peptides according to their precursor ion intensity; viral peptides are shown by colored circles. Sequences are shown for the top five most abundant viral peptides. Lines show the position of the two most abundant peptides in each nested set. **C.** HLA-ABC eluted viral peptides. A schematic representation of each source protein and the location of the eluted sequence is shown (first and last residues indicated). **D.** HLA-DR and HLA-DP eluted viral peptides. A schematic representation of each source protein with the location of each eluted sequence is shown (first and last residues indicated); the predicted core epitope in each sequence is underlined. Nested sets of eluted peptides comprising length variants with the same core epitope are shown by lines below the sequence. The peptide sequence highlighted in red was used for biochemical and immunological assays (see Table 1). In C and D, each eluted sequence or nested set was identified by “P” followed by a number. **E.** Label-free quantification of proteins present in infected cells; proteins were ranked from most to least abundant, with viral proteins highlighted in color. **F.** Relationship between viral protein abundance and eluted peptide abundance. For each source protein, the sum of intensities of all eluted peptides derived from it was used to calculate the peptide abundance.

**Table 1.**
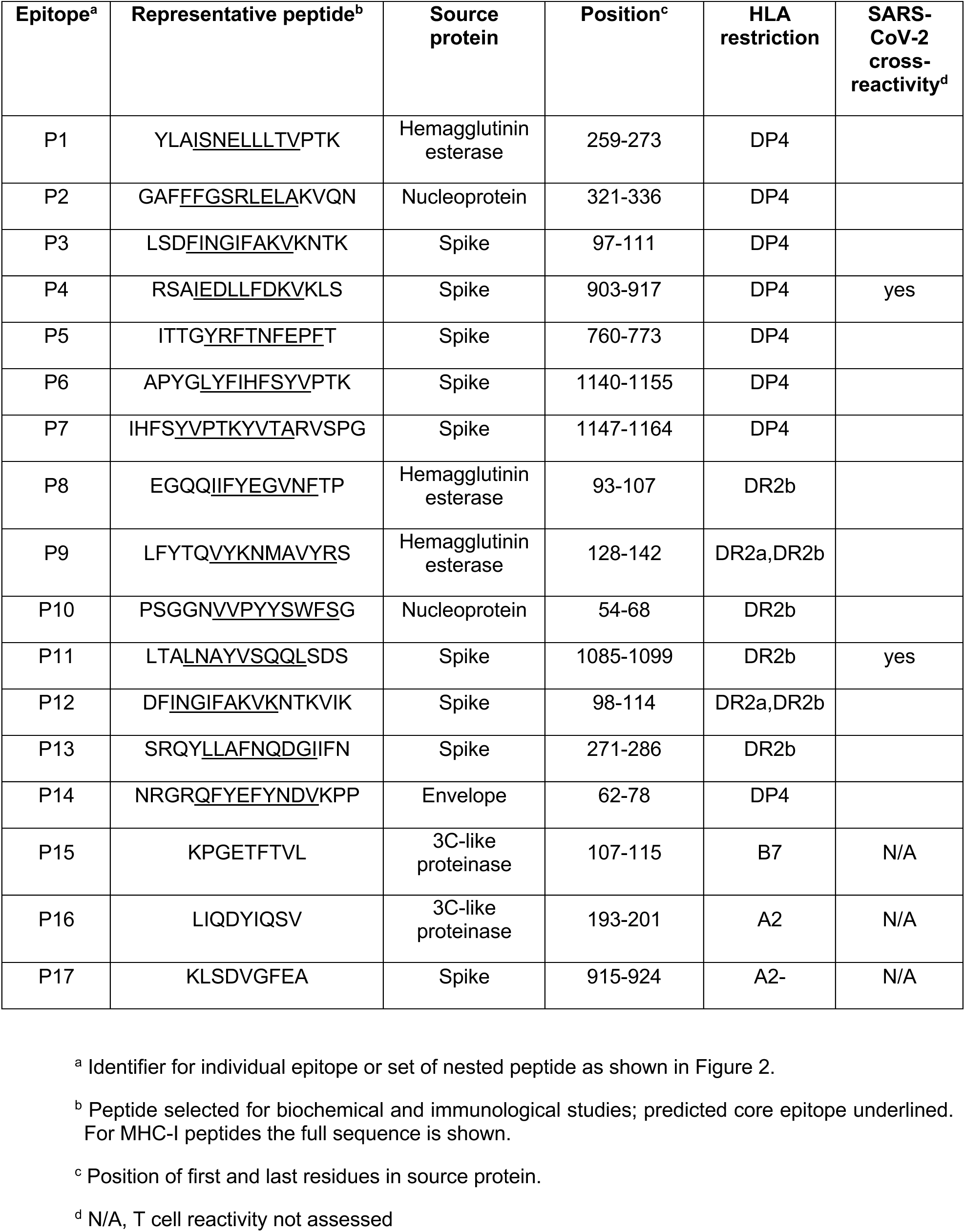
Naturally processed OC43 peptides.

The three MHC-I-binding viral peptides were identified at relatively low abundances within the overall MHC-I peptidome (Fig 2B, HLA-ABC, Table S2d). Peptide P17 (Fig 2C), derived from the spike protein, was assigned to HLA-A2 by motif analysis, with predicted binding in the top 0.5% (Table S2d). Peptides P15 and P16 (Fig 2C) were derived from the 3C-like proteinase of the ORF 1ab polyprotein and were assigned to HLA-B7 and HLA-A2 respectively, based on predicted binding within the top 0.5% for these alleles, although weak binding of peptide P16 to HLA-C7 was also predicted (1.5%-tile) (Table S2d). The low abundance of virus-derived peptides within the overall MHC-I peptidome might be a result of MHC-I immune-evasion mechanisms, similar to those reported for SARS-CoV-2 [45,55,56].

Eighty MHC-II-binding viral peptides were identified, derived from nucleoprotein, spike, hemagglutinin esterase (HE), and envelope proteins (Table S2d). Some of these were among the most abundant peptides identified in the MHC-II peptidomes: the most abundant peptide for HLA-DR and the third most abundant peptide for HLA-DP were virus-derived peptides (Fig 2B, HLA-DR and HLA-DP). Most of the MHC-II peptides were detected as part of nested sets of overlapping peptides, as characteristic of MHC-II peptidomes Fig 2D. The 35 HLA-DR peptides comprise five nested sets and one individual peptide (Fig 2D, P8-P13) and the 45 HLA-DP peptides comprise five nested sets and two individual peptides (Fig 2D, P1-P7, P14). The most abundant viral MHC-II peptides were derived from spike (P3, P4, P11) and nucleoprotein (P2, P10), with HE- and E-derived peptides present at lower abundance Fig 2B.

To relate the abundance of eluted peptides to the overall abundance of the source proteins, we performed proteomics analysis of intact proteins present in the infected cell lysate. Four viral proteins were detected: nucleoprotein, spike, HE, and the accessory protein N2. Label-free quantitative analysis showed that the most abundant protein was the nucleoprotein, followed by spike, and HE (Fig 2E). Spike, nucleoprotein, and HE proteins were also the major source proteins for the eluted peptides (Fig 2F).

### MHC-II allele restriction of eluted peptides

The nested sets of peptides characteristic of MHC-II peptidomes are comprised of length variants surrounding a 9-residue core epitope that includes the major sites of MHC-peptide interaction. This is believed to result from variable trimming of MHC-bound peptides by endosomal proteases, leaving different numbers of residues flanking the core regions. As expected, for each of the nested sets of peptides, the predicted core epitope (underlined in Fig 2D) was found in the center of the overlapping set. Core epitopes for the eluted peptides were among the top-ranked predicted binders for each protein (Fig S3A-C), helping to explain why these particular peptides were selected for presentation. For instance, the top-ranked predicted peptides for nucleoprotein, spike, and envelope contain the binding core from the HLA-DP-eluted peptides P2, P5, and P14, respectively (Fig S3A). Similarly, the top-ranked predicted peptides for nucleoprotein, spike, and HE contain the binding core from the HLA-DR-eluted peptides P10, P11, and P8, respectively (Fig S3B-C).

For HLA-DR, peptides were tentatively assigned to DR2a or DR2b by motif analysis. In some cases, one allele was clearly preferred, with predicted binding in the top 5th percentile to DR2b but not DR2a as for P8, P9, P10, and P11 peptides (Table S2d). P12 peptides were predicted to bind in the top 5^th^ percentile for both DR2b and DR2a, and P13 peptides were not predicted to bind to either DR2b or DR2a. For HLA-DP predicted binding was in the top 5^th^ percentile for P2, P3, P5, P6, P7, and P14 peptides, but P1 and P4 were below this threshold. To experimentally assess MHC-II peptide binding for the eluted peptides, we used a fluorescence polarization competition binding assay [57,58] with synthetic peptides and purified recombinant MHC proteins. For each set of nested peptides, we selected one abundant peptide containing the predicted binding core for the nested set and the allele of interest (Table S2d). These peptides are listed in Table 1. For DR2b, IC_50_ values were below 1 µM for all the HLA-DR-eluted peptides except P12, including P13 which was not predicted to bind (Fig S3D and Table S2d). For DR2a, IC_50_ values were below 1 µM for P12, as predicted, and also for P9. For DP4.1, only P1 and P5 of eight representative eluted peptides tested showed IC_50_ values below 1 µM, although all but P2 and P6 exhibited IC_50_ values below a more relaxed 10 µM criterion (Table S3).

### T cell recognition of eluted HLA-DR and HLA-DP viral peptides

We evaluated whether the naturally processed and presented viral peptides were recognized by circulating CD4 T cells in blood from healthy donors. We selected donors with a partial HLA match to HEK293 cells (donors expressing DRB1*15:01 and DRB5*01:01 for DR peptides, and donors expressing DPA1*01:03/DPB1*04:02 or DPB1*04:01 for DP peptides, Table S4). We expected prior exposure of these donors to OC43 or other seasonal coronaviruses, but serum was not available from these donors to confirm exposure serologically. We first assessed T cell responses directly ex vivo in PBMC samples using ELISpot assays with the same set of peptides as tested for MHC-II binding. Ex-vivo IFN-γ responses were measured in donors expressing at least one of the alleles of interest, by stimulating PBMCs with a pool of all DP or all DR peptides (Fig 3A). Positive responses were observed in most donors tested (6/9 for DP and 8/9 for DR). Responding T cells were present at low frequencies, which varied considerably between donors (0.007-0.057% for DP; 0.001-0.011% for DR). Note that in this assay other HLA alleles are present in the donors besides the HEK293 alleles used for the elution studies, but with very few exceptions these alleles are the best predicted binders among the HLA-DR, HLA-DP, and HLA-DQ alleles present in each donor (Table S5).

**Figure 3:**
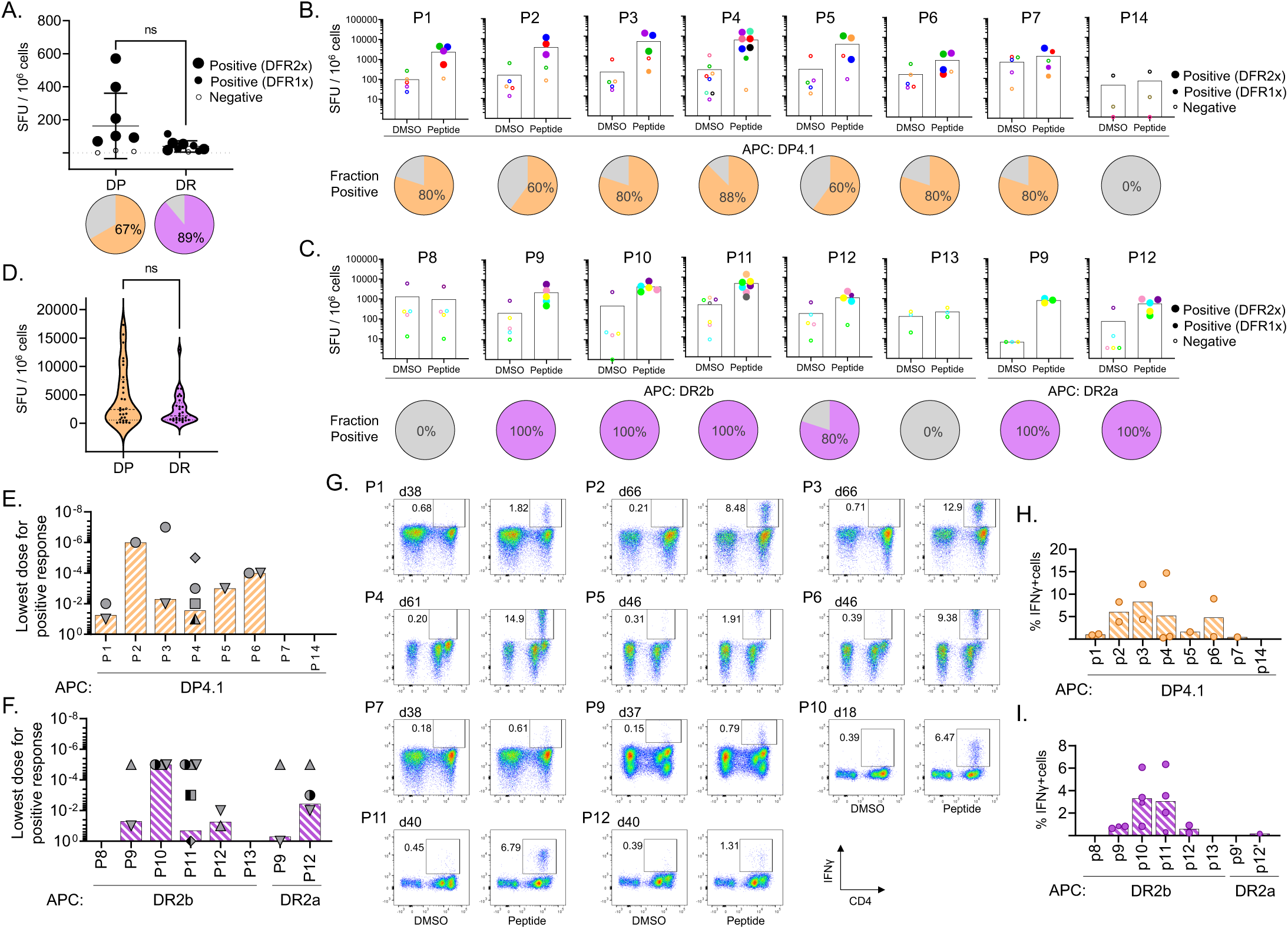
T cell recognition of eluted HLA-DR and HLA-DP viral peptides. **A.** Ex vivo T cell responses to OC43 eluted peptides (pooled by HLA allele) in pre-pandemic PBMC samples from donors with a partial HLA match to HEK293 cells. The plot shows IFN-γ production measured by ELISpot (SFU/10^6^ cells); pie graphs show the percentage of donors responding to the pool. **B-C.** Responding T cells from partially HLA-matched pre-pandemic donors were expanded in vitro by stimulation with each of the eluted peptides presented by a single allele antigen-presenting cells (APC). IFN-γ responses by expanded T cell populations from the same set of donors are shown in (B) for the HLA-DP peptides presented by DPA1*0301/DPB1*0401(DP4.1) and in (C) for the DR peptides presented by DRB1*1501 (DR2b) or DRB5*0101 (DR2a); pie graphs show the percentage of donors responding to the peptide. **D.** Summary of responses of single-peptide in vitro expanded T cells to the peptides, grouped by allele. **E-F.** Lowest peptide dose (10 – 10^-7^ µg/mL) eliciting a positive response to each eluted peptide, in experiments where the single-peptide in-vitro expanded T cells were tested for IFN-γ response to HLA-DP (E) or HLA-DR (F) eluted peptides presented by single allele APC (as in B-C). Each symbol represents a different donor. **G.** Response of single-peptide in-vitro expanded T cells to peptide stimulation followed in IFN-γ intracellular cytokine secretion (ICS) assay. Dot blots show CD4 expression (x-axis) and IFN-γ production (y-axis). DMSO, negative control. Responses > 3-fold background (DMSO) were considered positive. The gating strategy is presented in Figure S2. **H-I.** Summary of IFN-γ producing cell percentages in ICS assays for multiple donors for HLA-DP (H) and HLA-DR (I) peptides; only positive responses are shown. In A-C, statistical analysis to determine positive ELISpot responses was done by distribution-free resampling (DFR) method [89]; the size of the filled symbols indicates positive responses by DFR2x or DFR1x, while negative responses are shown as empty symbols. In A and D, statistical analysis was done by unpaired t-test (ns: not significant).

To increase the frequency of OC43-responding cells for detailed assessment of the responses to individual peptides, we expanded peptide-specific T cells in vitro. Using the expanded T cell populations, we measured IFN-γ production in response to re-stimulation with the same peptides, individually presented by single-allele antigen presenting cells (DPA1*01:03/DPB1*04:01 for P1-P7 and P14, DRB1*15:01 for P8-P13, and DRB5*01:01 for P9 and P12). Eleven peptides (all except P8, P13 and P14) showed individual positive responses by IFN-γ ELISpot in at least one of the donors analyzed (Fig 3B-C, bars and filled symbols), validating the presence of T cell responses to the peptide. Not every donor responded to every peptide, and different donors showed different patterns of responses. The fraction of donors who are positive for each of the responding peptides ranged from 60-100% (Fig 3B-C, pies). In general, responses were more frequently observed (p=0.006) in DR15 donors (80-100%) than in DP4 donors (60-88%), while responses were slightly stronger for DP peptides (3.7 ± 2.1×10^3^ SFU/10^6^ cells) than DR peptides (2.2 ± 1.8×10^3^ SFU/10^6^ cells) when tested at 1 µg/mL peptide concentration, although this difference is not significant (Fig 3D). There was a weak but significant correlation between the eluted peptide abundance (sum of precursor ion intensities by nested set) and the observed T cell response (r= 0.64, p= 0.009, Spearman). No correlation was observed between binding (predicted or experimental) and T cell responses, nor between binding and peptide abundance.

To explore the overall sensitivity of the different peptide-expanded T cells, dose-response experiments were performed, and the minimal activating peptide concentrations were determined (Fig 3E-F). In general, a wide range of minimal concentrations was observed. For instance, for P2 and P3 the minimal concentrations were 10^-6^ µg/mL and 10^-7^ µg/mL, respectively for expanded cells from donor 61, while for P11 (donor 07) and P9 (donor 40), the minimal concentration was 1 µg/mL. This indicates that T cells responding to P2 and P3 in donor 61 were more sensitive to lower peptide concentrations and may be able to respond more efficiently to infection. Within donors, differences in minimal concentration were observed for different peptides, suggesting a heterogeneous population that responds to different antigens with different efficiencies. In some cases, different donors showed similar sensitivity to a particular peptide, as is the case of P10 in donors 18, 22, and 40, which all responded at 10^-5^ µg/mL. However, in other cases, there was heterogeneity in the responses to a given peptide. For instance, for P4 the minimal concentration varied between 0.1 and 10^-5^ µg/mL in 4 donors. All these results may reflect the different history of exposure to OC43 and other coronaviruses and the evolution of the responding T cell repertoire in each individual, which translates to a lack of a clear hierarchy of functional avidity and immunodominance for most of the eluted peptides.

To characterize the T cells producing these responses, we performed intracellular cytokine staining (ICS) assays using the single-peptide-expanded T cell lines. As in the ELISpot assays, peptides P8, P13, and P14 did not produce a response. For the remaining 11 peptides, IFN-γ responses were observed exclusively in CD4+ T cell populations. Results from one representative cell line per peptide are shown in Fig 3G, with a summary of all results in Fig 3H-I. For 9 of these peptides, we were able to measure CD107a mobilization along with IFN-γ production (Fig S2B), and production of low levels of TNF-*α* was observed for 1 peptide (Fig S2C). No IL-2 or IL-10 production was observed for any peptide (not shown). This suggests that the CD4 T cells responding to the eluted OC43 peptides could be polyfunctional and have cytotoxic potential.

Altogether, these results present clear evidence of CD4+ T cells that recognize and respond to OC43-derived, DR2b, DR2a, and DP4.1/4.2-presented peptides, confirming the immunogenicity of these peptides in natural settings, showing that some of these peptides may be recognized by T cells at very low antigen concentrations in some donors, and highlighting the complexity of these responses.

### T cell cross-reactivity between OC43 and other human coronaviruses

The substantial sequence homology between OC43 and the other HCoVs (Fig S4A) raises the question of whether responding T cells could cross-react between the different orthologs. Sequence alignments of the naturally processed OC43 peptides with homologous sequences from other HCoVs are shown in Fig S4B, and a heatmap of conservation indices is shown in Fig S4C. Overall, the highest conservation is between OC43- and HKU1-derived peptides, with less for the other beta-coronaviruses MERS-CoV, SARS-CoV, and SARS-CoV-2, and even less for the alpha-coronaviruses 229E and NL63. Among the eluted peptides, P4, P6, and P11 are the most conserved across the 7 viruses and would be expected to have a high potential for cross-reactivity. The remaining peptides (P1, P2, P3, P5, P7, P8, P9, P10, P12, P13, and P14) were less conserved. Note that the HE protein, the source of the P1, P8, and P9 epitopes, is expressed by OC43 and HKU1 but does not have a homolog in any other HCoVs [7].

To evaluate experimentally the potential for cross-reactivity we initially focused on OC43 and SARS-CoV-2. We measured responses to the eluted OC43 peptides and their SARS-CoV-2 homologs, using T cell populations expanded with individual OC43 peptides from PBMC samples banked pre-pandemic before the outbreak of SARS-CoV-2 into the human population. Peptides with no homolog in SARS-CoV-2 (P1, P8, P9), or with no response in our donor pool (P8, P13, P14) were excluded. We measured T cell responses in single-peptide-expanded T cell lines using IFN-γ ELISpot assays, using partial-HLA-matched donors as before. Only the P4 and P11 SARS-CoV-2 homologs induced cross-reactive T cell responses in the single-peptide expanded lines (Fig 4A). Across a larger set of donors, similar cross-reactive responses were observed, with somewhat lower responses to the heterologous SARS-CoV-2 homologs than the OC43 peptides used for expansion (average 2-fold, p=0.044 for P4 and average 3.5-fold, p=0.011 for P11; paired t-test; Fig 4B). This indicates that a substantial proportion of T cells responding to the OC43-P4 and OC43-P11 peptides can cross-react with their SARS-CoV-2 homologs. To evaluate the sensitivity of these T cell lines to cross-reactive stimulation, we measured the dose-response to cognate and heterologous peptides. Robust cross-reactivity to heterologous stimulation was observed across the dose-response range for both P4 and P11 homologs in all donors tested, including pre-pandemic (Fig 4C) and those with recent COVID-19 infection (Fig 4D), with minimal stimulatory peptide concentrations in a wide range but similar for OC43 and SARS-CoV-2 homologs (Fig 4E).

**Figure 4:**
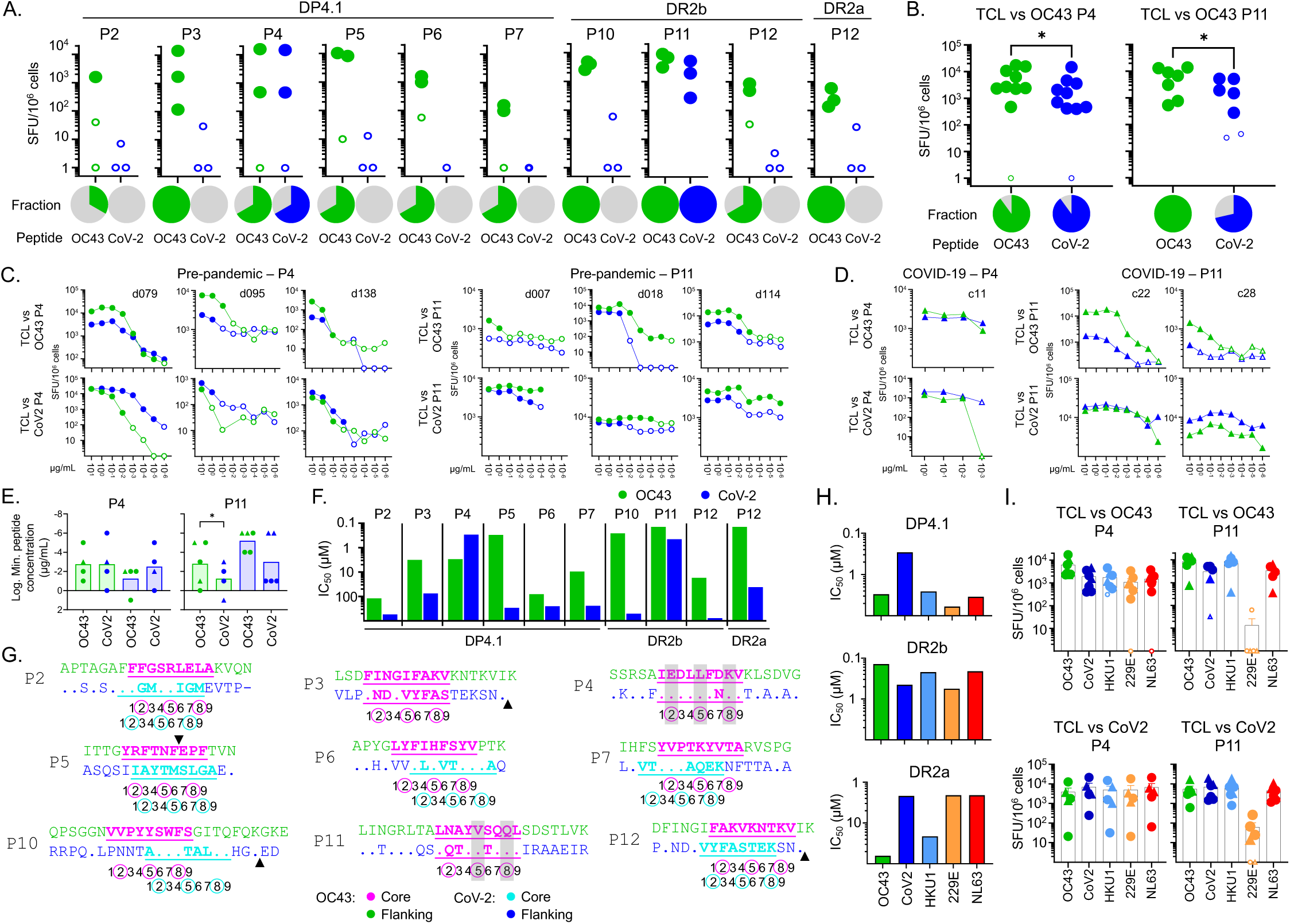
Epitope-specific T cell cross-reactivity between OC43 and other human coronaviruses. **A.** Screening of cross-reactive T cell responses in partially HLA-matched pre-pandemic donors. IFN-γ responses (SFU/10^6^ cells) to OC43 (green) or SARS-CoV-2 (blue) peptides using T cell lines expanded in vitro by stimulation with the eluted OC43 peptides and single allele APC. Pies show the fraction of responding donors to each peptide. **B.** For the two cross-reactive peptides (P4, P11), the screening was extended to more donors. **C.** Dose-response assay for the two cross-reactive peptides (P4, P11) in pre-pandemic donors. T cells were expanded in vitro with the OC43 peptide (TCL vs OC43, top row) or SARS-CoV-2 peptide (TCL vs CoV2, bottom row) and IFN-γ responses of each line to the OC43 peptide (green) or SARS-CoV-2 peptide (blue) were tested using single allele APC as before. **D.** Same as C but for COVID-19 convalescent donors. **E.** Lowest observed dose for a positive response for the cross-reactive peptides (tested in panels C and D). Pre-pandemic donors shown as circles and COVID-19 donors as triangles. **F.** Experimental binding of OC43 peptides (green) and the SARS-CoV-2 homologs (blue) to the relevant alleles. Half-maximal inhibitory concentration (IC_50_) values are shown. **G.** Sequence alignment of OC43 peptides and their SARS-CoV-2 homologs. OC43 sequences shown on top, with predicted core epitope shown in magenta and flanking regions in green; SARS-CoV-2 sequences on bottom, with residues different from OC43 shown and dots indicating identical residues. Predicted SARS-CoV-2 core epitope highlighted in turquoise with flanking regions shown in blue. Positions within the 9mer core epitope are indicated by numbers shown below the sequences; major T cell contacts are enclosed in circles. Arrowheads indicated gaps in the aligned sequences. If OC43 and SARS-CoV-2 epitopes are different both are shown. Gray bars show positions of identical residues at T cell contacts positions. **H.** Experimental binding of P4 and P11 OC43 peptides and their homologs in other coronaviruses to the relevant alleles. **I.** IFN-γ responses of T cell lines expanded in vitro with OC43 peptides (TCL vs OC43, top row) or with SARS-CoV-2 peptides (TCL vs CoV2, bottom row), to P4 and P11 peptides from OC43, SARS-CoV-2, and the other seasonal coronaviruses, presented by relevant single allele APC. In A-D and I, ELISpot statistical analysis by DFR method [89]; positive responses shown as filled symbols and negative responses as empty symbols. In B and E, statistical analysis was done by unpaired t-test. * p<0.05).

To explore factors that could have resulted in the observed pattern of OC43 and SARS-CoV-2 cross-reactive responses, we measured MHC binding of the SARS-CoV-2 homologs and compared them to the OC43 peptides (Fig 4F). We found weaker binding for most of the SARS-CoV-2 homologs, with the exception of P4, for which DP4.1 binding was 10-fold greater for the SARS-CoV-2 homolog. In addition to altering MHC binding affinity, amino acid substitutions can cause shifting of the preferred binding register, which would interfere with T cell recognition of homologous peptides. Of the nine peptides tested, only P3, P4, and P11 retain the predicted binding register in the SARS-CoV-2 homologs (Fig 4G), and only for P4 and P11 are the predicted T cell contacts completely or mostly conserved (shaded in Fig 4G).

We extended this analysis to the other seasonal human coronaviruses, using the T cell lines expanded in vitro with P4 and P11 peptides from pre-pandemic and COVID-19 donors. The P4 and P11 homologs from the seasonal coronaviruses mostly retained binding to DP4.1 (for P4) and DR2a/DR2b (for P11) (Fig 4H), and we measured the cross-reactive T response to these peptides. In general, all the P4- and P11-expanded T cell lines recognized each of the homologs, with the exception of P11 from 229E, which was recognized poorly by T cell lines expanded with SARS-CoV-2 or OC-43 homologs (Fig 4I).

## Discussion

The immune response to seasonal human coronaviruses is largely understudied and few T cell epitopes have been identified, although interest in this area has increased with the COVID-19 pandemic. To help fill this gap we identified naturally processed and presented viral epitopes expressed in OC43-infected cells using immunoaffinity purification of MHC-peptide complexes followed by mass spectrometry of eluted peptides. Only three viral peptides presented by MHC-I molecules were identified within the overall immunopeptidome of CIITA-transfected OC43-infected HEK293 cells, possibly due to virus-induced down-regulation of MHC-I expression. A total of 83 viral peptides presented by MHC-II molecules were identified, representing 14 distinct core epitopes present in nested sets characteristic of MHC-II processing. Eleven of these OC43-derived epitopes were recognized by recall responses in partially-HLA-matched donors. Almost all of the OC43-derived MHC epitopes identified in this work are reported here for the first time, although T responses to the two highly-cross-reactive epitopes P4 and P11 have been reported previously in studies characterizing seasonal coronavirus cross-reactivity to identified SARS-CoV-2 epitopes [21,25,27,28,59].

We identified only a few OC43-derived peptides presented by MHC-I molecules, and these were present at very low abundance within the overall MHC-I peptidome. One peptide from the spike protein and one from the 3C-like proteinase encoded by the ORF1ab polyprotein, both likely presented by HLA-A2, and a second 3C-like proteinase peptide likely presented by HLA-B7, were observed. These epitopes have not been previously reported, although a different OC43 spike epitope presented by HLA-24 [60,61] and two OC43-derived epitopes from other ORF1ab-derived proteins, both presented by HLA-A2 [62], have been described in studies of SARS-CoV-2 cross-reactive CD8 T cell responses. We observed potent MHC-I down-regulation after OC43 infection, which may have limited presentation of viral epitopes on MHC-I molecules. MHC-I down-regulation has not been previously reported for OC43, but is a common feature of many viruses [44–46], including SARS-CoV-2 [45,46,56,63]. Current understanding of SARS-CoV-2-induced MHC-I down-regulation points to a complex mechanism, with the involvement of several gene products: ORF3a reduces global trafficking of proteins including MHC-I [45], ORF6 inhibits induction of MHC-I by targeting the STAT1-IRF1-NLRC5 axis [63], ORF7a reduces cell-surface expression of MHC-I [45,46] by acting as *β*2-microglobulin mimic to interact with MHC-I heavy chain and slow its egress through the endoplasmic reticulum [45], and ORF8 also has been reported to down-regulate surface MHC-I through a direct interaction, although the specific mechanism is unclear [56]. However, none of these SARS-COV-2 gene products have significant homology with OC43, and elucidating the mechanism by which OC43 down-regulates MHC-I expression will require further investigation.

By contrast, eighty OC43-derived peptides presented by MHC-II molecules were found at high abundance within the overall MHC-II peptidome. Indeed, three of the top four most intense ions in the HLA-DR peptidome mass spectrum, and the third and fourth most intense ions in the HLA-DP peptidome mass spectrum, correspond to OC43-derived peptides. Most of the OC43-derived MHC-II-bound peptides were from spike and nucleoprotein, the major coronavirus structural proteins, consistent with the over-representation of these proteins we observed in the whole-cell proteome of infected cells. Several peptides derived from the hemagglutinin-esterase protein, which is believed to be required for cleavage of sialic acid residues to promote the release of progeny virus from infected cells, similarly to hemagglutinin-esterase proteins from influenza C and certain toroviruses and orthomyxoviruses [64]. Finally, one set of low-abundance peptides is derived from the small envelope protein. All the OC43-derived MHC-II-bound peptides were found as nested sets, except for three very low abundance peptides found as singletons. In each case, the nested sets surrounded the predicted nine-residue core epitope, with 1-9 residue extensions, consistent with endosomal protease trimming of MHC-bound peptides as expected in the MHC-II antigen-presentation pathway. We selected one representative peptide from each nested set to confirm binding to MHC-II, and to assign presenting MHC allotypes to the HLA-DR peptides, which could derive from either DR2a (DRB5*01:01) or DR2b (DRB1*05:01), both of which are expressed by HEK293 cells and co-purified with the LB3.1 antibody that we used for immunoaffinity. Each of the eight representative HLA-DP eluted peptides bound to DP4.1, although with varying affinity not entirely predicted by NetMHCIIpan4.1. Of the six representative HLA-DR peptides, one (P12) bound exclusively to DR2a, four exclusively to DR2b, and one to both allotypes (P9). As previously observed in another study of naturally processed MHC-II peptides in virus-infected cells [34], the eluted peptides generally were among the top predicted binders for each viral protein, one exception being P1 from the hemagglutinin-esterase protein.

We tested representative eluted peptides for recognition by T cells from HLA-matched donors. Of fourteen peptides tested, we observed robust T cell responses to eleven. In other systems, characterization of naturally-processed, MHC-bound peptides by mass spectrometry of infected cells has proven to be an efficient route for T cell epitope discovery [31,32,32–39,65,66]. We observed a correlation between the observed T cell response and epitope abundance in the overall immunopeptidome, whereas a significant correlation was not observed for the predicted or even observed peptide binding affinity. Thus, characterization of naturally-processed peptides from virus-infected cells can be a highly efficient epitope discovery approach, particularly compared to screening comprehensive overlapping peptides libraries or large sets of predicted MHC binders, where typically T cell responses are observed to only a small fraction of the candidate epitopes. A similar trend relating T cell response to epitope abundance has been observed in some [39] but not all [32,65,66] previous studies, although it should be noted that all of these previous studies involved CD8 T cell responses. Three eluted peptides (P8, P14, and P13) were not recognized by T cells from HLA-matched donors. These peptides were present at relatively low abundance in the peptidomes, although in some cases (P6, P7, P9) peptides with even lower abundance were recognized. We examined whether these peptides might not be immunogenic because of homology to self-peptides [67]. The peptides that were not recognized had similar homology scores to the closest matching self-peptides as did peptides that were recognized, although the number of exact matches in the core epitope region was somewhat larger for peptides that were not recognized (mean 6.3 vs 4.6, p=0.016).

Among human and animal coronaviruses, the approach of characterizing naturally-processed peptides presented by MHC proteins in infected cells to date has only been applied to SARS-CoV-2 [38,68]. Weingarten-Gabbay et al [38] eluted MHC-I bound peptides from SARS-CoV-2-infected A549 and HEK293 cell lines, and identified 28 canonical epitopes from spike, nucleoprotein, membrane, ORF7a, and several Orf1ab-derived nonstructural proteins, together with 9 non-conventional epitopes derived from out-of-frame transcripts in spike and nucleoprotein. Nagler et al [68] similarly identified two MHC-I epitopes derived from out-of-frame viral transcripts together with 11 conventional epitopes from spike, nucleoprotein, NSP1, and NSP3. We searched for such out-of-frame peptides in the OC43-derived immunopeptidome but did not find convincing evidence (see methods). As an alternative to infection, Pan et al [69] transfected cell lines with membrane or NSP13 genes and identified five MHC-I epitopes. In addition to the infection studies mentioned above, Nagler et al [68] also characterized MHC-bound peptides derived from cell lines transfected with individual nucleoprotein, envelope, membrane, and nsp6 genes, and identified additional MHC-I and also HLA-DR epitopes. Using a somewhat different experimental approach, Knierman et al [70] and Parker et al [71] added purified recombinant SARS-CoV-2 spike protein to monocyte-derived dendritic cells, which might simulate physiological antigen uptake by professional antigen-presenting cells at sites of infection. Peptides containing SARS-CoV-2 homologs of the OC43 P4 and P11 epitopes that we characterized here were among the many MHC-II-bound peptides that derived from the added recombinant proteins [70,71].

Several previous studies of the T cell response to SARS-CoV-2 in pre-pandemic donors have identified T cell responses that are cross-reactive with homologous epitopes from seasonal coronaviruses including OC43 [16,21,26,27,59,72–75]. However, there is still not a consensus on the involvement of the cross-reactive response in the clinical outcome, although recent studies have pointed to a role for cross-reactive CD8 T cell responses in protection from SARS-CoV-2 infection [62] and severe COVID-19 [76]. To identify additional cross-reactive epitopes, we tested the reactivity of T cell lines expanded with the eluted OC43 peptides for cross-reactivity with SARS-CoV2 homologs. Among the nine naturally processed CD4 T cell epitopes that were robustly recognized by donors in our cohorts, only two (P4 S_903-917_ and P11 S_1085-1099_) were targeted by T cells cross-reactive with SARS-CoV-2. Dose-response curves were similar for both SARS-CoV-2 and OC43 versions of the cross-reactive P4 and P11 epitopes, in both pre-pandemic and COVID-19 donors. This suggests that T cells might respond similarly during infections with either virus. Notably, these same epitopes were observed previously in an unbiased screen of SARS-CoV-2-derived peptides targeted by HCoV cross-reactive T cells [25], as well as in other studies of T cell responses cross-reactivity between SARS-CoV-2 and HCoVs [21,26–28,77–80]. For both the P4 and P11 epitopes, the OC43 and SARS-CoV-2 homologs are predicted to bind to the respective MHC-II proteins using the same binding frame, and peptide residues at the predicted T cell contact positions are identical or conserved. For the seven OC43-derived naturally processed T cell epitopes with SARS-CoV-2 homologs that were not targeted by cross-reactive responses, six had predicted shifts of the MHC-II binding frame caused by peptide substitutions at MHC-II contact positions. The one epitope for which the predicted MHC-II binding frame was preserved (P3 S_97-111_) has substitutions at each of the TCR contact positions, which would be expected to abrogate cross-reactive T cell binding. Thus, the pattern of observed CD4 T cell cross-reactivity can be explained by a simple model in which the key parameters are the preservation of the MHC-II binding frame and conservation of T cell receptor contact residues.

For studies of the differential response to SARS-CoV-2 and seasonal coronaviruses, epitopes specific to the seasonal coronaviruses are required. Among the OC43-eluted peptides for which cross-reactive T cell responses to SARS-CoV-2 homologs were not observed, P10 N_54-68_ elicited recall responses in all donors tested. Responding CD4 T cells showed a high sensitivity, with minimal peptide concentrations of about 10 pg/mL. This epitope is not strongly conserved among the HCoVs (Suppl Fig S4) and may be a good candidate to study and follow OC43-specific responses. In addition, epitopes P1 HE_259-273_, and P9 HE_128-142_ both are recognized by strong responses in a large majority of donors tested. human coronaviruses, only OC43 and HKU-1 express HE proteins, consistent with their use of 9-O-acetylated sialic acids as an entry receptor. Neither SARS-CoV-2 nor MERS-CoV, SARS-CoV, 229E, or NL63 express HE homologs. No HE-derived T cell epitopes have been reported from any other organism (although neutralizing antibodies to influenza C HE have been reported [81,82]. Thus, T cell responses to P1 HE_259-273_ and P9 HE_128-142_ would be expected to mark specific exposure to HCoVs (OC43 and/or HKU1) and might be useful in evaluating the contribution of HCoV exposure in SARS-CoV-2 incidence or pathogenesis.

There are some limitations to this study. The HEK cells used for immunopeptidome characterization were manipulated to ensure stable expression of MHC-II proteins by introducing the CIITA gene, which may favor the processing and presentation in the MHC-II compartment. In addition, these cells may not be representative of the natural targets of infection in the respiratory tract. Also, we assumed that the pre-pandemic donors would have been exposed to OC43. We did not consider T cell responses restricted by the mismatched MHC molecules. Finally, T cell responses not associated with IFN-γ, not able to expand with peptide stimulation in vitro, or below our detection level would have been missed by our approach.

In summary, we characterized the spectrum of naturally-processed viral peptides presented by MHC molecules in HEK293.CIITA cells infected with the human seasonal coronavirus OC43. MHC-II presented peptides dominated the OC43-derived viral immunopeptidome, possibly due to the potent down-regulation of MHC-I molecules in infected cells. The spike protein is the major source of OC43-derived epitopes, with contributions from nucleoprotein and hemagglutinin-esterase. Most of the naturally-processed peptides are recognized by T cells from HLA-matched donors. Three seasonal-coronavirus-specific CD4 T cell epitopes and two SARS-CoV-2-cross-reactive CD4 epitopes were identified. These epitopes provide a basis for studies of the cellular immune response to OC43, and for evaluating the role of pre-existing seasonal coronavirus immunity in SARS-CoV-2 infection and vaccination.

## Materials and Methods

### Cell lines

HEK293 cells were kindly provided by Dr. Kenneth Rock (UMass Chan Medical School). Cells were maintained in DMEM medium supplemented with L-glutamine (2 mM), sodium pyruvate (1 mM), non-essential amino acids (1 mM), and 10%FBS 37°C/5% CO_2_. HEK293 cells were transduced using the LentiORF® clone of CIITA (OriGene RC222253L3). The cells were selected using puromycin selection marker for 2 passages over the period of 7 days. The cells were further transduced using human *ace2* containing lentiviral particles, a kind gift from Dr. Rene Mehr (UMass Chan Medical School), to facilitate future work with other coronaviruses. The cells were stained for anti-HLA-DR, HLA-DP and HLA-DQ to confirm the MHC-II expression. These cells were further enriched by flow-based sorting for ACE2 expression and HLA-DR expression.

DP4.1-transfected cell line (M12C3, DPA1*0103/DPB1*0401, Williams et al., 2018) was kindly provided by Dr. S. Kent (UMass Chan Medical School). Cells were maintained in RPMI 1640 medium supplemented with L-glutamine (2 mM), penicillin (100 U/mL), streptomycin (100 mg/mL) and 10% FBS at 37°C/5% CO_2_.

Single HLA class II-transfected cell lines L466.2 (derived from the DRB1*15:01 cell line L466.1) and L416.3 (DRB5*01:01) [83] were kindly provided by Dr. Cecilia Sofie Lindestam Arlehamn (La Jolla Institute for Immunology). Cells were maintained in RPMI supplemented with L-glutamine (2 mM), penicillin (100 U/mL), streptomycin (100 mg/mL), non-essential amino acids (1 mM), Sodium Pyruvate (1 mM), G418 (200 µg/mL), and 10% FBS at 37°C/5% CO_2_. Sodium butyrate (100 mg/mL, Sigma B5887) was added the day before harvest to induce MHC expression.

### Virus production and cell infection

Human coronavirus OC43 strain VR-759 was obtained from ATCC (beta-coronavirus-1, #VR-1558). The virus was propagated in the lung fibroblast cell line MRC-5 (ATCC# CCL-171) at a multiplicity of infection (MOI) of 0.01 and the virus was collected after 5 days. Virus stocks were titrated using a standard TCID_50_ assay. HEK293.CIITA cells were infected at a MOI of 0.1 for 3 days, at which time the cells were collected, washed with PBS, and the cell pellets were frozen at −80°C until use. Percentage of infected cells were measured by intracellular staining for the nucleoprotein (mouse anti-coronavirus OC43 nucleoprotein clone 542-70, Millipore).

### Isolation of MHC Class I and Class II bound peptides

Detergent-solubilized fractions isolated from OC43 infected HEK293.CIITA cells were used for elution experiments. Cells were suspended in ice-cold hypotonic buffer (10 mm Tris-HCl, pH 8.0, containing protease inhibitors) and lysed using bath sonicator (Misonix S-4000 Ultrasonic Liquid Processor) maintained at 4°C with the amplitude of 70. The sonication was done for 3 mins with a cycle of pulse for 20 secs followed by resting cells on ice for 10 secs. Unlysed cells, nuclei, cytoskeleton, and cell debris were removed by centrifuging the lysate at 2000 ×g for 5 min at 4 °C. The supernatant was collected and further centrifuged at 100,000 ×g for 1 h at 4 °C to pellet the membrane/microsome fraction. This fraction was solubilized in ice-cold 50 mM Tris-HCl, 150 mM NaCl, pH 8.0 and 5% β-octylglucoside in a dounce homogenizer and incubated on ice for 1 hour. Benzonase (50 U/mL), 2 mM MgCl_2_, and protease inhibitor cocktail, were added to inactivate virus, and the mixture was rotated slowly overnight at 4 °C. Solubilized membranes were centrifuged at 100,000 ×g for 1 hour at 4 °C and the supernatant used for MHC-peptide isolation and immunopeptidome characterization. The supernatant was equilibrated with protein A agarose beads and isotype antibody conjugated beads sequentially for 1 hour each at 4 °C and allowed to mix slowly to remove nonspecific binding proteins. The precleared membrane fraction was then incubated sequentially with immunoaffinity beads of protein A agarose-LB3.1 antibody (HLA-DR), protein A agarose-B7/21 antibody (HLA-DP), and protein A agarose-W6/32 (HLA-ABC) antibody sequentially for 2 hours each at 4 °C and allowed to mix slowly. The beads were washed with several buffers in succession as follows: (1) 50 mM Tris-HCl, 150 mM NaCl, pH 8.0, containing protease inhibitors and 5% β-octylglucoside (5 times the bead volume); (2) 50 mM Tris-HCl, 150 mM NaCl, pH 8.0, containing protease inhibitors and 1% β-octylglucoside (10 times the bead volume); (3) 50 mM Tris-HCl, 150 mM NaCl, pH 8.0, containing protease inhibitors (30 times the bead volume); (4) 50 mM Tris-HCl, 300 mM NaCl, pH 8.0, containing protease inhibitors (10 times the bead volume); (5) PBS (30 times the bead volume); and (6) HPLC water (100 times the bead volume). Bound complexes were acid-eluted using 2% TFA. Detergent, buffer components, and MHC proteins were removed using a Vydac C18 microspin column (The Nest Group, Ipswich, MA). The mixture of MHC and peptides were bound to the column, and after washes with 0.1% TFA, the peptides were eluted using 30% acetonitrile in 0.1% TFA. Eluted peptides were lyophilized using a SpeedVac and were resuspended in 25 µL of 5% acetonitrile and 0.1% TFA.

### Liquid Chromatography–Mass Spectrometry (MS)

For LC/MS/MS analysis, peptide extracts were reconstituted in 7 µL of 5% acetonitrile containing 0.1% (v/v) trifluoroacetic acid and separated on a nanoACQUITY (Waters Corporation, Milford, MA). A 3.5 µL injection was loaded in 5% acetonitrile containing 0.1% formic acid at 4.0 µL/min for 4.0 min onto a 100 µm I.D. fused-silica precolumn packed with 2 cm of 5 µm (200 Å) Magic C18AQ (Bruker-Michrom, Auburn, CA) and eluted using a gradient at 300 nL/min onto a 75 µm I.D. analytical column packed with 25 cm of 3 µm (100 Å) Magic C18AQ particles to a gravity-pulled tip. The solvents were A) water (0.1% formic acid); and B) acetonitrile (0.1% formic acid). A linear gradient was developed from 5% solvent A to 35% solvent B in 60 min. Ions were introduced by positive electrospray ionization via liquid junction into a Orbitrap Fusion™ Lumos™ Tribrid™ Mass Spectrometer. Mass spectra were acquired over m/z 300–1,750 at 70,000 resolution (m/z-200), and data-dependent acquisition selected the top 10 most abundant precursor ions in each scan for tandem mass spectrometry by HCD fragmentation using an isolation width of 1.6 Da, collision energy of 27, and a resolution of 17,500.

### Peptide Identification

Raw data files were peak processed with Proteome Discoverer (version 2.1, Thermo Fisher Scientific) prior to database searching with Mascot Server (version 2.5, Matrix Science, Boston, MA) against a combined database of UniProt_Human, UniProt_hCoV-OC43 and an out-of-frame OC43 unconventional ORF database constructed according to Stern-Ginossar et al [84]. Search parameters included no-enzyme specificity to detect peptides generated by cleavage after any residue. The variable modifications of oxidized methionine and pyroglutamic acid for N-terminal glutamine were considered. The mass tolerances were 10 ppm for the precursor and 0.05 Da for the fragments. Search results were then loaded into the Scaffold Viewer (Proteome Software, Inc., Portland, OR) for peptide/protein validation and label-free quantitation. Scaffold assigns probabilities using PeptideProphet or the LDFR algorithm for peptide identification and the ProteinProphet algorithm for protein identification, allowing the peptide and protein identification to be scored on the level of probability. An estimated FDR of 5% was achieved by adjusting peptide identification probability. Peptides identified in a blank run were excluded from the peptidomes. Peptides with Mascot Ion score below 15 were also excluded. Only one match to the OC43 unconventional ORF database was identified for an HLA-DP-bound peptide. This sequence (LTILYLWVGIILSVIVL), derived from an out-of-frame ORF in the membrane gene, did not match the HLA-DP binding motif and the single-ion spectrum was poor, so this sequence was not considered further.

### Label-free proteomic analysis

Flow-through samples from the affinity columns used for immunopeptidome studies were collected and used for label-free proteomics analysis studies. 1 µg of the flow-through was trypsin digested using S-Trap™ Mini Spin Column (PROTIFI). An injection of ∼200 ng was loaded by a Waters nanoACQUITY UPLC in 5% acetonitrile (0.1% formic acid) at 4.0 μl/min for 4.0 min onto a 100 μm I.D. fused-silica precolumn packed with 2 cm of 5 μm (200 Å) Magic C18AQ (Bruker-Michrom). Peptides were eluted at 300 nL/min from a 75 μm I.D. gravity-pulled analytical column packed with 25 cm of 3 μm (100 Å) Magic C18AQ particles using a linear gradient from 5–35% of mobile phase B (acetonitrile + 0.1% formic acid) in mobile phase A (water + 0.1% formic acid) for 120 min. Ions were introduced by positive electrospray ionization via liquid junction at 1.5kV into a Orbitrap Fusion™ Lumos™ Tribrid™ Mass Spectrometer. Mass spectra were acquired over m/z 300–1,750 at 70,000 resolution (m/z 200) with an AGC target of 1e6, and data-dependent acquisition selected the top 10 most abundant precursor ions for tandem mass spectrometry by HCD fragmentation using an isolation width of 1.6 Da, max fill time of 110 ms, and AGC target of 1e5. Peptides were fragmented by a normalized collisional energy of 27, and fragment spectra acquired at a resolution of 17,500 (m/z 200). Raw data files were peak-processed with Proteome Discoverer (version 1.4, Thermo Scientific) followed by identification using Mascot Server (version 2.5, Matrix Science) against an UniProt_Human, UniProt_hCoV-OC43 and out-of-frame hCoV-OC43 databases. Search parameters included Trypsin/P specificity, up to 2 missed cleavages, a minimum of two peptides, a fixed modification of carbamidomethyl cysteine, and variable modifications of oxidized methionine, pyroglutamic acid for Q, and N-terminal acetylation. Assignments were made using a 10-ppm mass tolerance for the precursor and 0.05 Da mass tolerance for the fragments. All nonfiltered search results were processed by Scaffold (version 4.4.4, Proteome Software, Inc.) utilizing the Trans-Proteomic Pipeline (Institute for Systems Biology) with a 1% false-discovery rate. The data was processed using MaxQuant as well which uses Andromeda search engine and search parameters were kept the same as Mascot Server. The search was performed against a concatenated target-decoy database with modified reversing of protein sequences. For MHC protein quantitation, HLA-ABC heavy (alpha) chains and HLA-DR, HLA-DQ, HLA-DP beta chains were considered. Intensities of HLA-DRB1*15:01 and HLA-DRB1*01:01 were summed to provide an HLA-DR value. Peptides unique to HLA-C or HLA-E, a non-classical class I MHC bound by W6/32 along with HLA-ABC [85], were not detected, although two peptides identical in HLA-C and HLA-A were detected and assigned to HLA-A, and two peptides identical in HLA-E and HLA-B were detected and assigned to HLA-B.

### Gibbs Clustering

GibbsCluster-2.0 [86] within DTU Health Tech server, was used to align the eluted peptide sequences and analyze the motifs, which were displayed with Seq2Logo 2.0 [87]. We allowed the software to include cluster sizes of 1-5 with a motif length of 9 amino acids and clustering sequence weighting. Default values were used for other parameters: number of seeds =1, penalty factor for inter-cluster similarity =0.8, small cluster weight =5, no outlier removal, iterations per temperature step =10, Monte Carlo temperature =1.5, intervals for indel, single peptide and phase-shift moves = 10, 20, and 100, respectively, and Uniprot amino acid frequencies were used. For each sample, we selected the cluster that included the largest number of peptides analyzed. For HLA-DR and HLA-DP peptides, a preference for hydrophobic residue at P1 was used to align the motifs at the P1 position. For HLA-ABC peptides, MHC-I ligands of length 8-13 residues parameters were loaded. The fraction of sequences that contributed to each cluster is shown in the figures.

### Peptide binding assay

We used a fluorescence polarization competition binding assay, modified from one developed for MHC-I peptide binding [57], to measure peptide binding affinity to soluble recombinant MHC-II molecules. Soluble DRB1*15:01 and DRB5*01:01 with a covalently linked CLIP peptides [88] were a gift of Drs. John Altman and Richard Willis (Emory University and NIH Tetramer Core Facility). Soluble DP4 (HLA-DPA1∗01:03/DPB1∗04:01) with a covalently-linked CLIP peptide was prepared essentially as described [88]. Human oxytocinase EKKYFAATQFEPLAARL, MBP peptide NPVVHFFKNIVTPR and influenza hemagglutinin PRFVKQNTLRLAT peptide were labeled with Alexa Fluor 488 (Alexa488) tetrafluorophenyl ester (Invitrogen, Carlsbad, CA) and used as probe peptides for DP4.1, DR2b and DR2a binding. Binding reactions were carried out at 37°C in 100 mM sodium citrate, 50 mM sodium chloride, 0.1% octyl β-D-glucopyranoside, 5 mM ethylenediaminetetraacetic acid, 0.1% sodium azide, 0.2 mM iodoacetic acid, 1 mM dithiothreitol as described [58] for peptide-free HLA-DR1, but with 1 U/µg thrombin (DP4.1) or 3C protease (DR2b and DR2a) added to cleave the CLIP linker and HLA-DM included to initiate peptide exchange. Thrombin or 3C protease enzymes was inactivated after 3 hours of reaction using protease cocktail inhibitor, and the reaction was continued for 24 hours at 37 °C before FP measurement using a Victor X5 Multilabel plate reader (PerkinElmer, Shelton, CT). DP4.1-Clip (250 nM), DR2b-CLIP (500 nM) and DR2a-CLIP (250 nM) concentrations were selected to provide 50% maximum binding of 25 nM probe peptide in the presence of 500 nM soluble HLA-DM. Binding reactions also contained serial dilutions of test peptides with 5-fold dilutions. The capacity of each test peptide to compete for binding of probe peptide was measured by the fluorescence polarization (FP) after 24 hours at 37 °C. FP values were converted to fraction bound by calculating [(FP_sample - FP_free)/(FP_no_comp - FP_free)], where FP_sample represents the FP value in the presence of test peptide; FP_free represents the value for free Alexa488-conjugated respective peptide; and FP_no_comp represents values in the absence of competitor peptide. We plotted fraction bound versus concentration of test peptide and fit the curve to the equation *y* = 1/(1 + [pep]/IC_50_), where [pep] is the concentration of test peptide, *y* is the fraction of probe peptide bound at that concentration of test peptide, and IC_50_ is the 50% inhibitory concentration of the test peptide.

### ELISpot assay

IFN-γ ELISpots were performed using Human IFN gamma ELISpot KIT (Invitrogen, San Diego, CA) and MultiScreen Immobilon-P 96 well filtration plates (EMD Millipore, Burlington, MA), following the manufacturer’s instructions. Assays were performed in CST^TM^ OpTmizer^TM^ T cell medium (Gibco, Grand Island, NY). Peptides or peptides pools were used at a final concentration of 1 µg/mL per peptide (10 - 10^-7^ µg/mL for dose-responses curves); as negative controls were used DMSO (DMSO, Fisher Scientific, Hampton, NH) and a pool of human self-peptides (Self-1 [34]), and PHA-M (Gibco, Grand Island, NY) was used as a positive control. For ex vivo assays, PBMC were incubated with peptides or controls for ∼48 hours. We used 4×10^5^ cells per well. For assays with cells expanded in vitro, ∼5×10^4^ cells per well were incubated with an equal number of irradiated single allele APCs in the presence of peptides or controls for ∼18 hours. Two to four wells of each peptide, pool of peptides, or PHA-M, and at least 6 wells for DMSO were usually tested. Secreted IFN-γ was detected following the manufacturer’s protocol. Plates were analyzed using the CTL ImmunoSpot Image Analyzer (ImmunoSpot, Cleveland, OH) and ImmunoSpot 7 software. Statistical analysis to determine positive responses was performed using the distribution-free resampling (DFR) method described by Moodie et al [89].

### Intracellular cytokine secretion assay (ICS)

ICS was performed using in vitro expanded T cells as previously described [34] with minor modifications. Briefly, single allele APCs were resuspended in CRPMI (w/o phenol red) +10% fetal bovine serum (FBS, R&D Systems) containing 1 µg/mL of each peptide and incubated overnight. On the day of the assay, T cell lines were collected, washed, and resuspended in the same medium and added to the pulsed APCs (1:1 ratio); at this time, anti-CD107a-CF594 was added, followed by the addition of brefeldin A and monesin at the suggested concentrations (Golgi plug / Golgi stop, BD Biosciences, San Jose, CA). After 6 hours of incubation, cells were collected, washed, and stained using a standard protocol, which included: staining for dead cells with Live/Dead Fixable Aqua Dead Cell Stain Kit^TM^ (Life Technologies, Thermo Fisher Scientific, Waltham, MA); blocking of Fc receptors with human Ig (Sigma-Aldrich, St. Louis, MO); surface staining with mouse anti-human CD3-APC-H7, CD4-PerCPCy5.5, CD8-APC-R700, CD14-BV510, CD19-BV510, CD56-BV510; fixation and permeabilization using BD Cytofix/Cytoperm^TM^; and intracellular staining with mouse anti-human IFN-γ-V450, TNF-α-PE-Cy7, IL-2-BV650, (all from BD Biosciences, San Jose, CA). Data were acquired using a BD LRSII flow cytometer equipped with BD FACSDiva software (BD Biosciences, San Jose, CA) and analyzed using FlowJo v.10.7 (FlowJo, LLC, Ashland, OR). The gating strategy consisted in selecting lymphocytes and single cells, followed by discarding cells in the dump channel (dead, CD14+, CD19+, and CD56+ cells), and selecting CD3+ cells in the resulting population.

### Peptides and HLA binding predictions

Peptides for these studies were obtained from 21^st^ Century Biochemicals (Marlborough, MA) and BEI Resources (Manassas, VA). Peptide sequences using in the assays are shown in Table S6. HLA-peptide binding prediction was performed with NetMHCpan4.1 or NetMHCIIpan4.0 (Reynisson et al., 2020) for peptides eluted from MHC-I and MHC-II proteins, respectively. Sequence logo of predicted motifs obtained using Motif Viewer in NetMHCpan or NetMHCIIpan. The Immune Epitope Database IEDB [19] was used to search for T cell responses to seasonal and pandemic coronavirus epitopes.

### Sequence conservation analysis

We selected one representative strain from each human coronavirus: OC43 strain VR759 (NC_006213), HKU1 Isolate N1 (NC_006577), Human beta-coronavirus 2c EMC/2012 (JX869059), SARS coronavirus Tor2 (NC_004718), SARS-CoV-2/human/USA/WA-CDC-02982585-001/2020 (MT020880), Human coronavirus 229E (AF304460), and NL63 strain Amsterdam I (NC_005831).Sequence alignment of spike, nucleoprotein, hemagglutinin esterase, and envelope proteins were generated using Clustal Omega v1.2.4 [90]. Conservation indices for each position of the alignment were calculated using the AL2CO algorithm [91] using the alignment previously generated and the default settings. Human Peptides sequences Eluted peptides sequences were searched against the whole human proteome to find potential human homologs

## Acknowledgements

The authors wish to thank Dr. John Altman and Dr. Richard Willis (Emory University) for DRB1*15:01 and DRB5:01:01 proteins used for peptide binding studies, Liying Lu for antibodies used for immunoaffinity purification, Dr. Rene Mehr for ACE2 lentivirus, and Dr. Kenneth Rock for HEK293 cells. We acknowledge the assistance of Nadia Sultana and the UMass Chan Medical School Flow Cytometery and Mass Spectrometry Facilities.

## Data Availability

LC-MS/MS data have been deposited to the ProteomeXchange Consortium via the MassIVE repository with the dataset identifier MSV000090595. To access the files, use login web access as MSV000090595_reviewer. All other relevant data are within the manuscript and its Supporting Information files

## Author Contributions

ABA: Conceptualization, Formal Analysis, Investigation (virology, T cell studies, cross-reactivity analysis), Methodology, Validation, Visualization, Writing – original draft preparation, Writing-review & editing

PPN: Conceptualization, Formal Analysis, Investigation (HEK293/CIITA/Ace3 generation, immunopeptidome analysis, label-free proteomic analysis, binding studies), Methodology, Validation, Visualization, Supervision, Writing-original draft preparation, Writing-review & editing.

JMCC: Conceptualization, Data Curation, Formal Analysis, Investigation (T cell studies, cross-reactivity analysis), Methodology, Supervision, Validation, Writing – original draft preparation, Writing-review & editing

MK: Methodology (immunopeptidomics)

GCW: Investigation (protein biochemistry)

SAS: Methodology (mass spectrometry), Writing – Review and Editing

LJS: Funding Acquisition, Conceptualization, Methodology, Project Administration, Supervision Visualization, Writing – original draft preparation, Writing-review & editing.

## Supporting information

**Figure S1: MHC-I and MHC-II alleles present in HEK293 cells and sequence logos of the predicted 9mer core epitope**. From Motif Viewer within NetMHCpan 4.1 and NetMHCIIpan4.0 (DTU Health Tech).

**Figure S2: Polyfunctional response elicited by OC43 eluted peptides**. A. Gating strategy for ICS experiments. B. CD107a staining of single-peptide in vitro expanded T cells responses to the expanding peptide presented by single allele APC. Dot blots show CD4 (x-axis) and CD107a expression on surface (y-axis). C. ICS for TNF-α production by single-peptide in vitro expanded T cells responses to the expanding peptide presented by single allele APC. Dot blots show CD4 expression (x-axis) and TNF-α production (y-axis). Dot plots for DMSO and peptide are shown. Responses > 3-fold background (DMSO) signal was considered positive.

**Figure S3: Eluted peptides binding predictions and experimental binding**. Epitope prediction on whole viral proteins / allele combination were obtained from NetMHCIIpan and sorted by score. Peptides containing the predicted core of the eluted peptides are highlighted in each protein. A. predictions for DP4.1 and DP4.2; B. predictions for DR2b; C. Predictions for DR2a. D. Experimental binding of eluted peptides to relevant alleles; dark colors indicate strong binding.

**Figure S4: Homology between OC43 and other human coronaviruses**. A. Percentage identity of OC43 proteins vs homologous proteins in other human coronaviruses (http://imed.med.ucm.es/Tools/sias.html). B. Sequence alignment of the 14 OC43 eluted peptides to positional homologs in other human coronaviruses. Whole OC43 sequence (with core epitope underlined), and differences in the other sequences are shown. For each alignment, the conservation score at each position was obtained using AL2CO algorithm and presented as a bar graph, with the core epitope positions in black. C. Summary of conservation scores for each eluted peptide to each of their homolog peptides in other human coronaviruses. Scores normalized to 100% identity to OC43 peptide as 1, and no conservation as 0. An average per peptide is shown at the bottom of the heatmap. NA indicates no homolog protein between OC43 and the corresponding virus.

**Table S1:** Cellular proteomics analysis on the OC43 infected and uninfected HEK293.CIITA cells. S1a. Summary of host and viral protein identified in infected and/or uninfected cells. One biological replicate of uninfected cells and two biological replicates of OC43-infected cells were analyzed, with each having two technical replicates. Average and standard deviation of technical / biological replicates are presented in the table. S1b. MHC-I and MHC-II levels in uninfected and OC43-infected HEK293.CIITA cells as measured by proteomics quantitative analysis.

**Table S2:** Immunopeptidome of OC43-infected HEK293.CIITA cells. S2a. HLA-ABC immunopeptidome; S2b. HLA-DR immunopeptidome; S2c. HLA-DP immunopeptidome; S2d. OC43 immunopeptidome. For each peptide, mass spectrometry identification parameters are shown (eluted sequence, length, source protein, intensity, Scaffold identification probability, and Mascot Ion and Identity scores). In addition, NetMHCpan 4.1 or NetMHCIIpan 4.0 predictions were performed and predicted core for each relevant allele, score, and rank are shown for each peptide.

**Table S3:** Binding affinities of OC43 eluted peptides and homologs. S3a. Binding affinities of HLA-DR OC43 eluted peptides and homologs in other coronaviruses to DR2b and DR2a. S3b. Binding affinities of HLA-DP OC43 eluted peptides and homologs in other coronaviruses to DP4.1. In both tables: Binding affinities of OC43 eluted peptides and the corresponding homologs in other coronaviruses are shown as IC_50_ (µM) of each peptide to the indicated HLA. The mean and the standard deviation (SD) of two independent experiments are presented.

**Table S4:** Donors used in the study.

**Table S5:** Binding predictions of OC43 eluted peptides to other alleles present in the donors used in the study.

**Table S6:** Synthetic peptides used in the study.

